# What are the gray and white matter volumes of the human spinal cord?

**DOI:** 10.1101/2020.07.01.182444

**Authors:** Simon Henmar, Erik B. Simonsen, Rune W. Berg

## Abstract

The gray matter of the spinal cord is the seat of somata of various types of neurons devoted to the sensory and motor activities of the limbs and trunk as well as a part of the autonomic nervous system. The volume of the spinal gray matter is an indicator of the local neuronal processing and this can decrease due to atrophy associated with degenerative diseases and injury. Nevertheless, the absolute volume of the human spinal cord has rarely been reported, if ever. Here, we use high–resolution magnetic resonance imaging, with a cross–sectional resolution of 50 × 50*μ*m^2^ and a voxel size of 0.0005mm^3^, to estimate the total gray and white matter volume of a post mortem human female spinal cord. Segregation of gray and white matter was accomplished using deep learning image segmentation. Further, we include data from a male spinal cord of a previously published study. The gray and white matter volumes were found to be 2.87 and 11.33 ml, respectively for the female and 3.55 and 19.33 ml, respectively for a male. The gray and white matter profiles along the vertebral axis were found to be strikingly similar and the volumes of the cervical, thoracic and lumbosacral sections were almost equal.

**NEW AND NOTEWORTHY:** Here, we combine high field MRI (9.4T) and deep learning for a post-mortem reconstruction of the gray and white matter in human spinal cords. We report a minuscule total gray matter volume of 2.87 ml for a female and 3.55 ml for a male. For comparison, these volumes correspond approximately to the distal digit of the little finger.

## Introduction

The gray and white matter of the central nervous system are conspicuous properties that are attractive to quantify and compare across humans and animal species. These features represent fundamental properties of the nervous system: the gray matter is the territory of the input-output processing units, i.e. the neuronal somata, and the white matter consists primarily of the myelin tracks that connect them. The gray and white matter (GM and WM) volume also serve as a diagnostic metric in neurological conditions such as multiple sclerosis where the volume decreases due to axonal atrophy (Kearney et al., 2015; Gilmore et al., 2005; Schmierer and Miquel, 2018; Pallebage-Gamarallage et al., 2018; Losseff et al., 1996). The gray matter can also decrease due to atrophy of neurodegenerative diseases such as dementia and in particular in the spinal cord where amyotrophic lateral sclerosis causes motor neuronal death and decline in the gray matter volume (Paquin et al., 2018). Spinal abnormalities such as cysts and syringomyelia can induce pressure om the GM and WM and reduce the effective volume of healthy tissue surrounding the syrinx. Similarly, the extent of damage from spinal cord injury from fall, motor vehicle accident or ischemia can be assessed via the cross-sectional area of GM and WM (Seif et al., 2018; Kakulas, 2004). Hence, the GM and WM cross-sectional areas serve important diagnostic metric for spinal neurology (Papinutto et al., 2015; Klein et al., 2011). A convenient technique for measuring the GM and WM areas is via the non-invasive magnetic resonance imaging (MRI), and quantitative MRI has been increasingly helpful for diagnosis and measuring the integrity in such conditions.

Although the cross-sectional areas and partial volumes of GM and WM and the ratio between them have been reported previously, e.g. (Tsagkas et al., 2019; Wheeler-Kingshott et al., 2014), their total volume have rarely been reported, if ever. Here, we measure the total volume of the GM and WM in the spinal cord of a healthy human female using high-resolution MRI in post mortem condition. The post mortem condition permitted long continuous scanning (approximately 30 hrs) and the absence of movement combined with a strong magnetic field (9.4T) enabled high image quality and a fine cross-sectional resolution (50×50*μ*m^2^) with a slice thickness of 200*μ*m (voxel size of 0.0005mm^3^), hence facilitating a precise estimate of the volumes. With such high resolution and large number of slices (n=1919), manual segmentation of gray and white matter is unfeasible and automated segmentation is required. Automated segmentation via artificial intelligence has previously been implemented to accomplish such a task and we employed that method for the identification of gray and white matter zones (Perone et al., 2018). Data from a male subject was provided by a previously published study on a post mortem spinal cord, which was used for a different purpose (Calabrese et al., 2018). The scanning details of this study was slightly different and although the male was reportedly healthy, their post morten analysis revealed a small lesion at the 6th thoracic nerve level.

## Methods

The study was conducted on a deceased individual who had bequeathed her body to science and education at the Department of Cellular and Molecular Medicine (ICMM) of Copenhagen University according to Danish legislation (Health Law No 546, Section 188). The study was approved by the head of the Body Donation Program at ICMM. This study consists of high-resolution MRI scanning of two human spinal cords, one based on publicly available data set of a male (please find details in (Calabrese et al., 2018)), the other a female performed in this study. A spinal cord from a 91-year old Caucasian female without known diseases was dissected out and fixed using immersion fixation in paraformaldehyde (4%) within 24 hours after death. The tissue was kept in fixative for 2 weeks, after which it was transferred to and stored in phosphate buffered saline.

### Structural MRI acquisition

Images from the female spinal cord were acquired on a 9.4 T preclinical MRI system (BioSpec 94/30; Bruker Biospin, Ettlingen, Germany) equipped with a 1.5 T/m gradient coil. Prior to imaging, the spinal cord was placed in a plexiglass tube and immersed in fluorinert (FC-40 Sigma Aldrich) to reduce background signal. Because of the limited field of view (1.6 cm) relative to the length of the spinal cord (approximately 40 cm), scanning was done in 29 sections. Between each section scan, the spinal cord was advanced 1.4 cm by a custom-built mechanical stepper, resulting in a 0.2 cm overlap of neighboring sections. For each section, a T2-weighted 2D RARE structural scan was performed.

Scan parameters were TR=7 s, TE=30 ms, 20 averages, field of view 1.92 × 1.92 × 1.6 cm^3^, and a matrix size of 384 × 384 × 80, resulting in 50 × 50 *μ*m^2^ in-plane resolution and a slice thickness of 200 *μ*m, resulting in a voxel size of 500 000 *μ*m^3^. Scan time was approximately 30 hours.

## MRI data analysis

### Stitching

For each of the 29 scanned sections, dicom-files were exported from the scanner software (ParaVision 360, Bruker) and converted to nifti format using open source software (dcm2niix, (Li et al., 2016)). The individual section images were combined by rigid registration of overlapping regions using another piece of open source software (Advanced Normalization Tools Python adaption, ANTSpy, with default parameters (Avants et al., 2009)). To stitch two sections A and B together, the overlapping 0.2 cm bottom of A and top of B were registered to each other. A margin of displacement in the z-direction (rostral-caudal) of 400 *μ*m (two slices) was allowed. For each such displacement window, registration between overlapping regions was run 50 times. The window with the highest mean mutual information across the 50 runs was accepted. Section B was then transformed to match section A using the transformation matrix from the best registration among the 50 runs. The bottom of the transformed section B was then registered to the top of the next section in the same way as A and B were registered to each other. This process was repeated until the last section had been registered to the second last one. The transformed sections were then concatenated using half of the overlapping region from each section, resulting in a single image of the whole spinal cord.

### Cropping

The stitched image was cropped to reduce deep learning memory load. A mask separating spinal cord from background was created for the stitched image using an open source software (MRtrix’s dwi2mask) on a faked 4D version of the 3D image (Tournier et al., 2019). Cropping was done to accommodate the mask in-plane, with a 10 voxel (0.5 mm) margin. The caudal-most part of the spinal cord was removed from the image since it did not contain any internal GM or WM. Final cropped image dimensions were 348 × 296 × 1919 voxels, 17.4 × 14.8 × 383.8 mm^3^.

### Definition of spinal segments

Spinal segments were defined using the relative lengths of each spinal segment estimated from a number of studies on human spinal morphology (Frostell et al., 2016).

### Segmentation of gray and white matter

Segmentation of images into gray matter and white matter was done on the stitched image using a 2D convolutional neural network with dilated convolutions developed for GM segmentation in spinal MRI (Perone et al., 2018). Model code was adapted from the Spinal Cord Toolbox (De Leener et al., 2017) to enable training on our data. A training data set was constructed by manual labelling of 15 slices. The model was trained on these slices using data augmentation with details described elsewhere (Perone et al., 2018). Training parameters were batch size 9, 32 batches per epoch, and 734 epochs. This procedure was done separately for GM and WM. Voxels present in both masks were removed from the WM mask. Both masks were visually inspected and identified errors were manually corrected. The area of GM and WM was calculated for each slice as the number of voxels belonging to each of the masks multiplied by the in-plane resolution. For the male spinal cord, GM and WM masks published along with the structural image were used (Calabrese et al., 2018).

### Estimation of uncertainty in volume

The area of GM and WM was calculated based on the artificial intelligence image segmentation (Deep dilated convolutional neural network, (Perone et al., 2018)). The variability of area between neighboring slices is an indication of the certainty of the identification. Hence, we estimated the uncertainty in areas based on the mean and standard deviation of five consecutive slices sliding across the entire sample. The standard error of the mean was calculated and used as uncertainty and the propagation of uncertainty was estimated by summation for the total volumes and the fractions (Table 1). This was also performed on the adopted data (a male spinal cord) from a previously published study (Calabrese et al., 2018).

**TABLE 1.** Total volumes of gray and white matter from two human subjects calculated using high resolution MRI sequential scans and a deep dilated convolutional neural network for image segmentation of gray and white matter. † Calculated with permission from (Calabrese et al., 2018).

| Spinal cord volume |  |  |  |  |
| --- | --- | --- | --- | --- |
| Subject | Gray Matter (ml) | White matter (ml) | Total (ml) | Fraction (WM/GM) |
| Female 91-yr old | 2.87 ± 0.02 | 11.33 ± 0.03 | 14.20 ± 0.05 | 3.95 ± 0.04 |
| Male 60+ yrs old† | 3.55 ± 0.02 | 19.33 ± 0.03 | 22.88 ± 0.05 | 5.45 ± 0.04 |

## Results

First, the spinal cord from a 91-year-old female was analyzed. After training the machine learning algorithm (Perone et al., 2018) on selected slices, the gray and white matter was segregated and identified reliably as verified by visual inspection (Fig. 1A-B). In this specimen the first two cervical segments (C1-C2) were partially missing, making the cord a bit shorter. The cross-sectional area of the GM and WM masks was determined and plotted as a function of vertebral level (Fig. 1C). The GM and WM areas were larger both in the cervical and the lumbo-sacral enlargements. The fact that the white matter had a local peak in the lumbar enlargement suggests that some of the myelinated fibres both started and stopped within the lumbar region and thus serve local connectivity within the region. A similar local peak in the white matter profile was observed in the cervical region, which most likely represents the local connectivity within the cervical region.

**FIGURE 1.**
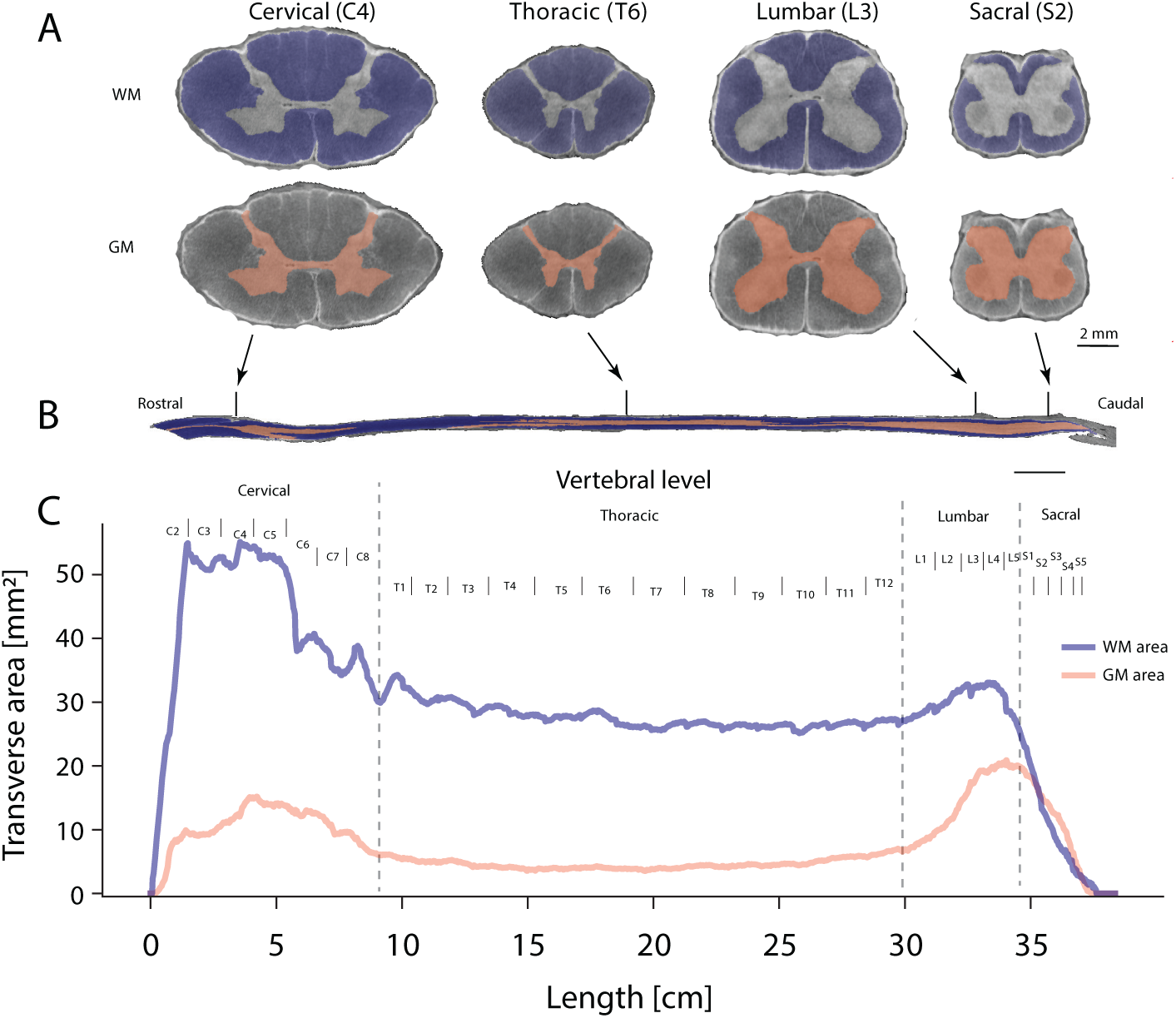
Estimates of gray and white matter area of a human spinal cord (91-year old female) as a function of vertebral level. *A:* Sample sections of cervical, thoracic, lumbar and sacral spinal cord from a T2-weighted MRI scan. Artificial intelligence identified the gray and white matter in each transverse slice (total of n=1919 slices), here shown with masks of white matter (blue, top) and gray matter (red, bottom). In-plane resolution is 50 × 50 *μ*m^2^. *B:* Location of samples indicated in a parasagittal section of the whole spinal cord. *C:* The cross-sectional GM and WM area as function of location. Vertebral levels are indicated.

Next, data from the spinal cord of a male in his sixties that was previously published for a different purpose (Calabrese et al., 2018) was analyzed in a similar fashion and compared with that from the 91-year-old female (Fig. 2). Since the length and size of the spinal cords are different presumably due to differences in their body size, we rescaled the profile of the the gray matter for vertebral levels (Fig. 2A). The profiles where strikingly similar except for a minor offset due to the missing initial cervical sections of the female spinal cord (C1-C2). An analogous comparison between the two subjects was performed for the white matter (Fig. 2B). Again, the profiles of the white matter along the vertebral axis were notably similar when the first subject (the female) data was rescaled to the other subject (same proportions as in Fig. 2A).

**FIGURE 2.**
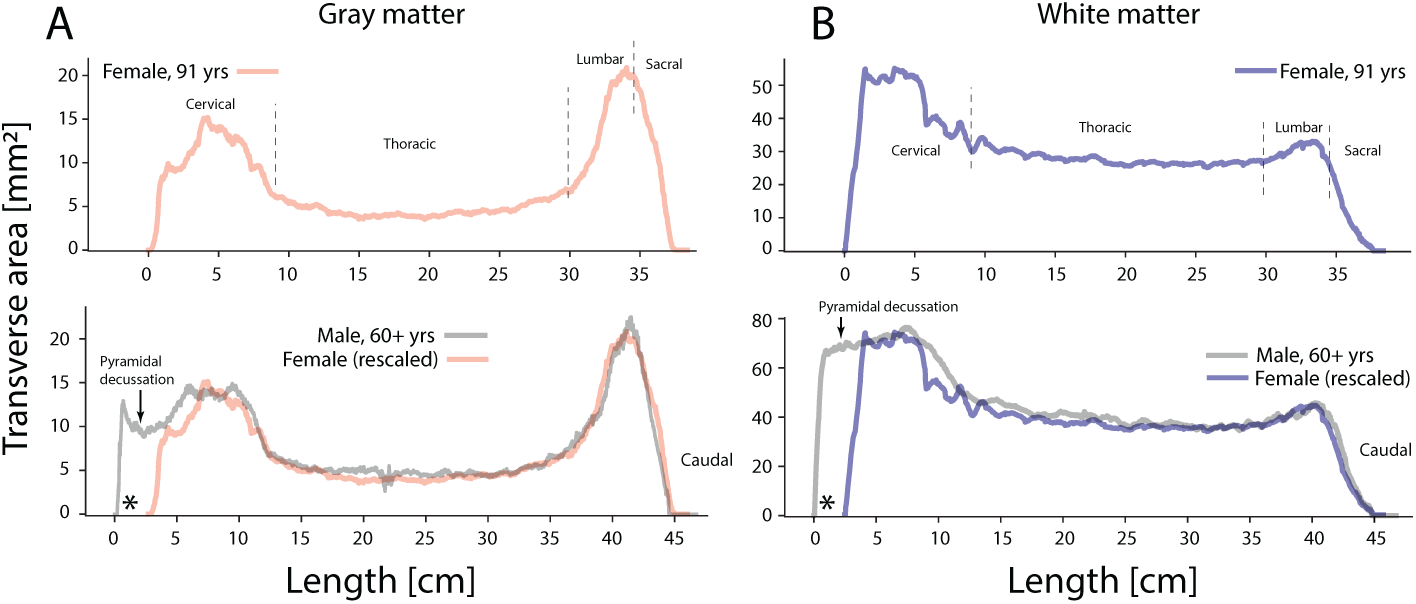
Comparison of profiles of spinal gray and white matter area along vertebral level of a female (91-year old) and a male (in his sixties). *A:* Gray matter of female (top) and a male (bottom, gray line). For comparison and due to difference in size, the female curve is rescaled and plotted together with the male profile (bottom). *B:* White matter vertebral profiles plotted and rescaled in similar fashion to (A). Note the female spinal cord was missing C1-C2, so the curves do not start at the same location (indicated by a *). Arrows indicate the pyramidal decussation. Male data adapted with permission from (Calabrese et al., 2018).

The total volume of gray and white matter was calculated by integrating the cross-sectional areas with the section thickness of 200 *μ*m (Table 1). The total gray matter volume was a minuscule 2.87 ml and 3.55 ml for the female and male, respectively. For the male data, the part of the gray matter profile more rostral of the pyramidal decussation was not included, since that is part of the caudal medulla oblongata. The white matter volume was approximately 4-5 times larger than the gray matter volumes (Table 1).

From the profiles (Fig. 2A) it is clear that the thoracic cross-sectional area is much smaller than the cervical and lumbar parts, but it is also much longer. Interestingly, when separating out the gray matter volumes in their cervical, thoracic and lumbosacral parts, these were approximately equal to each other (Fig. 3). For the female spinal cord data, the volumes were 0.92, 0.98 and 0.97 ml, for the cervical, thoracic and lumbosacral sections, respectively. Similar equivalence was found in the male specimen, although the cervical was slightly larger (Fig. 3). Hence, it seems that the extra length of the thoracic part makes up for the smaller cross-sectional area that it has in the vertebral profile (Fig. 2A). In contrast to the gray matter volumes, the white matter volumes were greatly different across the cervical, thoracic and lumbosacral regions (Fig. 3).

**FIGURE 3.**
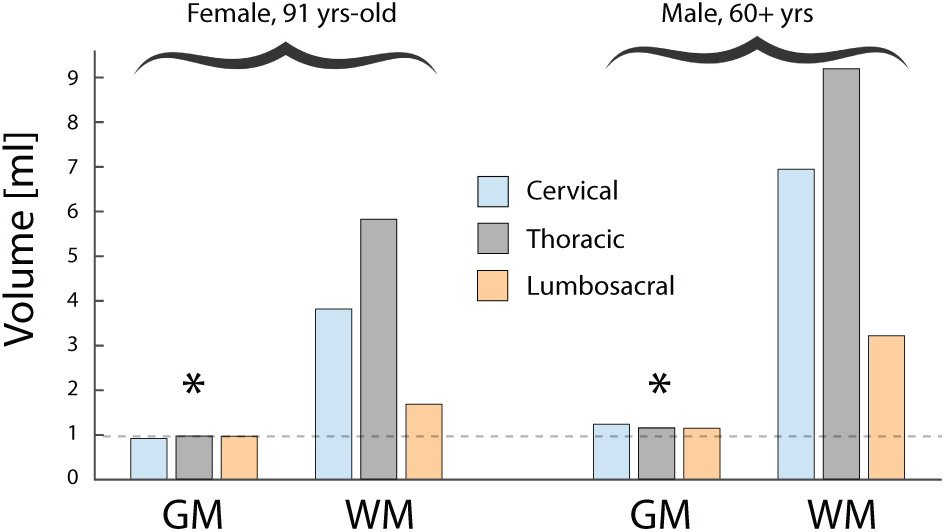
Gray and white matter volumes distributed over the cervical, thoracic and lumbosacral parts for the female and male subjects. There is a striking similarity in gray matter volumes across the sections (see *) despite their difference in length and cross-sectional area. In contrast, the white matter volumes are vastly different across the 3 sections. Male data adapted and analysed with permission from (Calabrese et al., 2018).

## Discussion

Gray and white matter are distinguished features of the central nervous system. Nevertheless, measurements of their volumes in the spinal cord have rarely been reported, possibly due to the laborious nature of assessment by conventional histological techniques and its long extension. Magnetic resonance imaging performed on living subjects is limited on field strength and scan time, and by various types of tissue and air in proximity of the tissue of interest, which distorts the MRI-signal. This restricts the image resolution and quality. The analysis of post-mortem tissue where scanning time is no restriction opens up new possibilities. The combination of high-strength magnetic fields in a smaller bore absent of mechanical movement artifacts and where the tissue, in a signal-free, non-distorting fluid, can be advanced through the smaller scanning region with strong gradients allows for unprecedented spatial resolutions (Atik et al., 2019; Sitek et al., 2019; Calabrese et al., 2018; Colon-Perez et al., 2015; Lundell et al., 2011). In the present study, we use a small bore strong field (9.4T) scanner where a post-mortem ex-vivo spinal cord is advanced using a stepping motor through a small region of about 2 cm of scanning area with constant field strength and homogeneous gradient. The sections from these scans formed the basis for estimating the gray and white matter volumes and the profiles of their cross-sectional areas.

Nevertheless, the benefit of in-vivo scanning, e.g. (Tsagkas et al., 2019; Papinutto et al., 2015; De Leener et al., 2018), is a much larger cohort which is devoid of any post-mortem distortion due to tissue processing and which allows better statistical sampling. The age of our subjects should also be considered, since the gray and white matter volumes change over the course of a lifetime (Kearney et al., 2015). Younger adults would be expected to have slightly larger volumes than reported here (Table 1). The two spinal cords were obtained post-mortem and dissected out within 24 hours. One was treated in paraformaldehyde (4% formaldehyde) and the other was fixed using formalin, which is a saturated formaldehyde solution, with the same fixative effect on the tissue. The difference in concentration should have minimal effect, if any, on the tissue properties. Similarly, the lesion in the thoracic region of the male specimen, which likely occurred post-mortem during extraction, did not have a clear effect on the cross-sectional profile (Fig. 2).

Two observations from the profiles (Fig. 2) are interesting. First, the cross-sectional area of the gray matter within the lumbosacral enlargement is significantly larger than of the cervical enlargement. Nevertheless, the cervical enlargement is longer and the volume of the two enlargements turns out to be almost identical (Fig. 3). If the gray matter volume represents local circuitry, which can be interpreted as computational capacity, then the sensorimotor requirements of the hands and the arms are approximately equivalent to those of the legs. The thoracic region has the lowest cross-sectional gray matter area, but it also entails the longest part of the spinal cord. Hence, the volume of these regions are almost identical, perhaps as a coincidence (Fig. 3). Second, the white matter has a local maximum both in the cervical and the lumbar enlargements. The fact that the white matter volume does not steadily increase towards the cerebrum suggest that some of the white matter fibers that are both initiated in the more caudal regions also terminate within the spinal cord in more rostral regions, although the fibers are probably bi-directional. These fibers, also called the proper fasciculi or spinospinal fasciculi, partially represent communication and coordination between sensorimotor regions of the spine.

## Acknowledgements

Thanks to J. Tranum for providing the spinal cord, P. Koch for building the mechanical stepper, and Y. Mori for assistance in the MRI scans. Thanks to Calabrese et al. (2018) for making their data publicly available. Funded by The Independent Research Fund, Denmark (R.W.B., S. H.), Carlsberg foundation, Lundbeck foundation and Maersk foundation (R.W.B.).

## Data sharing

The acquired data as well as the processed data and the segmentation masks are available as supplement with the publication. It is also available on the laboratory web site (http://berg-lab.net) or upon request to the corresponding author.

## Abbreviations

GM: gray matter
WM: white matter
MRI: magnetic resonance imaging.

